# Probing transcription factor combinatorics in different promoter classes and in enhancers

**DOI:** 10.1101/197418

**Authors:** Jimmy Vandel, Océane Cassan, Sophie Lèbre, Charles-Henri Lecellier, Laurent Bréhélin

## Abstract

In eukaryotic cells, transcription factors (TFs) are thought to act in a combinatorial way, by competing and collaborating to regulate common target genes. However, several questions remain regarding the conservation of these combinations among different gene classes, regulatory regions and cell types. We propose a new approach named TFcoop to infer the TF combinations involved in the binding of a target TF in a particular cell type. TFcoop aims to predict the binding sites of the target TF upon the binding affinity of all identified cooperating TFs. The set of cooperating TFs and model parameters are learned from ChIP-seq data of the target TF. We used TFcoop to investigate the TF combinations involved in the binding of 106 TFs on 41 cell types and in four regulatory regions: promoters of mRNAs, lncRNAs and pri-miRNAs, and enhancers. We first assess that TFcoop is accurate and outperforms simple PWM methods for predicting TF binding sites. Next, analysis of the learned models sheds light on important properties of TF combinations in different promoter classes and in enhancers. First, we show that combinations governing TF binding on enhancers are more cell-type specific than that governing binding in promoters. Second, for a given TF and cell type, we observe that TF combinations are different between promoters and en-hancers, but similar for promoters of mRNAs, lncRNAs and pri-miRNAs. Analysis of the TFs cooperating with the different targets show over-representation of pioneer TFs and a clear preference for TFs with binding motif composition similar to that of the target. Lastly, our models accurately distinguish promoters associated with specific biological processes.

## INTRODUCTION

Transcription factors (TFs) are regulatory proteins that bind DNA to activate or repress target gene transcription. TFs play a central role in controlling biological processes, and are often mis-regulated in diseases [1]. Technological developments over the last decade have allowed the characterization of binding preferences for many transcription factors both *in vitro* [2, 3] and *in vivo* [4]. The current view is that TF combinations underlie the specificity of eukaryotic gene expression regulation [5], with several TFs competing and collaborating to regulate common target genes. As reviewed in Morgunova et al. [6] and Reiter et al. [7], multiple mechanisms can lead to TF cooperation. In its simplest form, cooperation involves direct TF-TF interactions before any DNA binding. But cooperation can also be mediated through DNA, either with DNA providing additional stability to a TF-TF interaction [8], or even without any direct protein-protein interaction. Different mechanisms are possible for the later. For example, the binding of one TF may alter the DNA shape in a way that increases the binding affinity of another TF [6]. Another system is the pioneer/settler hierarchy described in Sherwood et al. [9], with settler TFs binding DNA only if adequate pioneer TFs have already bound to open the chromatin. Lastly, other authors have hypothesized a non-hierarchical cooperative system, with multiple concomitant TF bindings mediated by nucleosomes [10]. This is related to the “billboard” system proposed for enhancers [11]. On the other hand, TFs that belong to the same protein family usually share identical or similar motifs and may compete for sites that match both motifs [12].

Several papers have studied the combinatorics of TFs from a statistical point of view. Most works aim to identify co-occurring TF pairs, *i.e.* pair of TFs showing binding sites that are in closest proximity than one would expect by chance. These analyses have been done either on the basis of TF binding sites (TFBSs) predicted *in silico* [13, 14] or with TFBSs obtained from ChIP-seq experiments [15, 16]. Depending on the approach, different difficulties may arise. *In silico* predicted TFBSs are known to include large amount of false positives (see below), which may bias the analyses and impede the discovery of co-occurring TFBSs. On the other hand, studies based on ChIP-seq data require as many ChIP-seq data as the number of studied TFs, and hence are intrinsically limited by the availability of these data. Moreover, with hundreds (or even thousands) of sequences, a small co-occurrence tendency may be statistically significant, even if the effect is actually very weak and would not be biologically relevant. A few works have studied TF combinations in a more global way, above the TF pair level. For example, Teng et al. [17] have applied the “frequent itemset” methodology to identify sets of co-occurring TFBSs on the basis of ChIP-seq data. However, many questions remain on the molecular determinants orchestrating TF binding and combinations [18]. Notably, with the expanding coding capacity of the human genome [19, 20], it remains to determine whether the expression of all gene classes, in particular coding mRNAs, long non-coding(lnc)RNAs and micro(mi)RNAs, is controlled by similar TF combinations in a given cell type. Likewise, TFs control gene expression through the binding of promoters and enhancers, which harbor similar but also specific genomic features [21]. It is then not clear whether the binding preferences of a given TF are similar in enhancers and promoters.

Here, we analyze global TF combinations from a different perspective. Rather than identifying TF pairs/sets that co-occur more frequently than expected by chance, we aim to identify TF combinations that can be predictive of the binding of a target TF. More formally, given a class of regulatory sequences (for example 500bp around the TSSs of the coding genes) and a ChIP-seq experiment targeting a specific TF in a specific cell type, we aim to identify the combinations of TFs whose predicted TFBSs can be used for predicting which sequences are effectively bound by the target TF in this cell type. Hence, rather than using purely statistical co-occurrence analysis, we study TF combinations in the framework of a TFBS prediction problem. The approach has several advantages. First, a single ChIP-seq experiment is theoretically sufficient to identify all TFs cooperating/competing with the target TF in the target cell type. Next, if a TF is selected in the combination, this means that its predicted binding sites are indicative of the presence of the target TF, which limit the number of false positives and the problems of spurious statistical significances. Finally, the approach takes into account all TFs and can therefore identify all possible TF combinations not just TF pairs.

TFBSs are traditionally modeled with position weight matrices (PWMs) [22]. Several databases such as JASPAR [23], HOCOMOCO [24], and Transfac [25], propose position frequency matrices (PFM, which can be transformed in PWMs) for hundred of TFs. These PWMs can be used to scan sequences and identify TFBSs using tools such as FIMO [26] or MOODS [27]. However, while a PWM usually identifies thousands of potential binding sites for a given TF in the genome [28], ChIP-seq analyses have revealed that only a fraction of those sites are effectively bound [29]. There may be different reasons for this discrepancy between predictions and experiments. First, PWMs implicitly assume that the positions within a TFBS independently contribute to binding affinity. Several approaches have thus been proposed to account for positional dependencies within the TFBS (see for example [30, 31]). Other studies have focused on the TFBS genomic environment, revealing that TFs have a preferential nucleotide content in the flanking positions of their core binding sites [32, 33]. Beyond the primary nucleotide sequence, structural constraints may also affect TF binding. For example, it is thought that TFs use DNA shape features to distinguish binding sites with similar DNA sequences [34, 35]. Some attempts have thus been made to integrate DNA shapes information with PWMs [36, 37]. Other studies have investigated the link between TF binding and epigenetic marks, showing that many TFs bind regions associated with specific histone marks [38]. Similarly, ChIP-seq experiments also revealed that most TFBSs fall within highly accessible (*i.e.*, nucleosome-depleted) DNA regions [39]. Consequently, several studies have proposed to supplement PWM information with DNA accessibility data to identify the active TFBSs in a given cell type [40, 41, 42]. However, it remains unclear whether these chromatin states are a cause or a consequence of TF binding [43]. Hence, while these approaches may be very informative for predicting TF binding, they should be used with caution if the goal is also to identify the DNA determinants of the binding. Besides, these approaches do not take into account TF combinations, which, as already discussed, may be important determinants of TF binding. For this reason, studying TF combinations through a TFBS prediction problem appears as an appealing approach.

It is important to note that beyond approaches based on known PWMs, several *ab initio* methods have also been proposed recently for predicting TFBSs from raw data sequences. Notably, deep learning approaches based on neural networks have proved to give higher prediction accuracy than simple PWM-based methods [44, 45]. However, *ab initio* methods, and particularly neural network approaches, are difficult to interpret (the inherent trade-off between accuracy and interpretability). Although some attempts have been made to post-analyze learned neural networks (see for example [46]), studying TF combinations and DNA determinants of TF binding from these models is not straightforward.

Hence, we devised a simple non *ab initio* strategy names TFcoop that predicts if a target TF binds a sequence of interest using two kinds of variables: i) the binding affinity (*i.e.* PWM affinity score) of the target TF as well as any other TF identified as cooperating with the target TF; and ii) the nucleotide composition of the sequence. TFcoop is based on a logistic model. The set of cooperating TFs and the model parameters are learned from ChIP-seq data of the target TF via LASSO penalization [47]. Learning can be done using a moderate amount of data, which allows us to learn specific models for different types of regulatory sequences. Using ChIP-seq data from the ENCODE project, we applied TFcoop to investigate the TF combinations involved in the binding of 106 different TFs on 41 different cell types and in four different regulatory regions: promoters of mRNAs, lncRNAs and miRNAs, and enhancers [48, 19, 49, 20]. We first showed that the approach outperforms simple PWM methods and has surprisingly good accuracy, close to that of *ab initio* methods like DeepSea [44]. We next assessed with independent experimental data that the cooperative TFs predicted by TFcoop actually bind the same regulatory sequences as the target TF. Then, we used TFcoop to analyze TF combinations in different cell types and regulatory regions. First, we show that TF combinations governing the binding of the target TF on promoters are similar for different cell-types but distinct in the case of enhancer binding. Second, for a given TF, we observe that TF combinations are different between promoters and enhancers, but similar for promoters of all gene classes (mRNAs, lncRNAs, and miRNAs). Analysis of the composition of TFs cooperating with the different targets show over-representation of pioneer TFs [9], especially in promoters, as well as binding sites with nucleotide composition similar to that of the target TF. We also observed that cooperating TFs are enriched for TFs whose binding is weakened by methylation [50]. Lastly, our models can accurately distinguish promoters into classes associated with specific biological processes.

## RESULTS

### Computational approach

Given a target TF, the TFcoop method identifies the TFBS combination that is indicative of the TF presence in a regulatory region. We first considered the promoter region of all mRNAs (defined as the 1000bp centered around gene start). TFcoop is based on a logistic model that predicts the presence of the target TF in a particular promoter using two kinds of variables: PWM affinity scores and (di)nucleotide frequencies (see Material and methods). For each promoter sequence, we computed the affinity score of the 638 JASPAR PWMs (redundant vertebrate collection), and the frequency of every mono- and dinucleotide in the promoter. These variables were then used to train a logistic model that aims to predict the outcome of a particular ChIP-seq experiment in mRNA promoters. Namely, every promoter sequence with a ChIP-seq peak is considered as a positive example, while the other sequences are considered as negative examples (see below). In the experiments below, we used 409 ChIP-seq datasets from ENCODE and different models. Each model targets one TF and one cell type. Given a ChIP-seq experiment, the learning process involves selecting the PWMs and (di)nucleotides that can help discriminate between positive and negative sequences, and estimate the model parameters that minimize prediction error. Note that the learning algorithm can select any predictive variable including the PWM of the target TF. See Material and methods for more details on the data and logistic model.

We used two different procedures for selecting the positive and negative examples. Each procedure actually defines a different prediction problem. In the first case, we kept all positive sequences (*i.e.* promoters overlapping a ChIP-seq peak in the considered ChIP-seq experiment), and randomly selected the same number of negative sequences among all sequences that do not overlap a ChIP-seq peak. In the second case, we used an additional dataset that measures gene expression in the same cell type as the ChIP-seq data. We then selected all positive sequences with non zero expression level and randomly selected the same number of negative sequences among all sequences that do not overlap a ChIP-seq peak but that have a similar expression level as the selected positive sequences. Hence, in this case (hereafter called the expression-controlled case), we learn a model that predicts the binding of a target TF in a promoter knowing that the corresponding gene is expressed. On the contrary, in the first case we learn a model that predicts the binding without knowledge about gene expression. The purpose of the expression-controlled case is to decipher TF combinations independently of the effect of epigenetic modifications that are linked to expression (e.g. DNA methylation and various histone marks). As all selected sequences are associated with expressed genes, the positives and negatives sequences are likely to be associated with the same epigenetic marks.

### TFcoop assessment

We ran TFcoop on the 409 ChIP-seq datasets and for the two prediction problems. The accuracy of each model was assessed by cross-validation by measuring the area under the Receiver Operating Curve (ROC). For comparison, we also measured the accuracy of the classical approach that discriminates between positive and negative sequences using only the affinity score of the PWM associated with the target TF. In addition, we estimated the accuracy of the TRAP method, which uses a biophysically inspired model to compute PWM affinity [51] and that of the approach proposed in [36], which integrates DNA shape information with PWMs. As shown in Figure 1 and Supp. Figures 1 and 2, TFcoop outperforms these PWM-based approaches on many TFs. Note that these comparisons are rather unfavourable for our method because they integrate all 69 CTCF experiments, while TFcoop has similar accuracy than classical PWM methods on this TF (see Supp. Figures 2). Note also that we ran TFcoop with tri- and quadri-nucleotide frequencies in addition to di-nucleotide frequencies. Although a consistent AUC improvement was observed, the increase was very slight most of the time (Supp. Figure 3). Lastly, we compared TFcoop accuracy to that of the deep learning approach DeepSea [44] and observed very close results (see Figure 1(b)). Hence, TFcoop performances appear to be in the range of that of classical *ab initio* methods.

**FIGURE 1.**
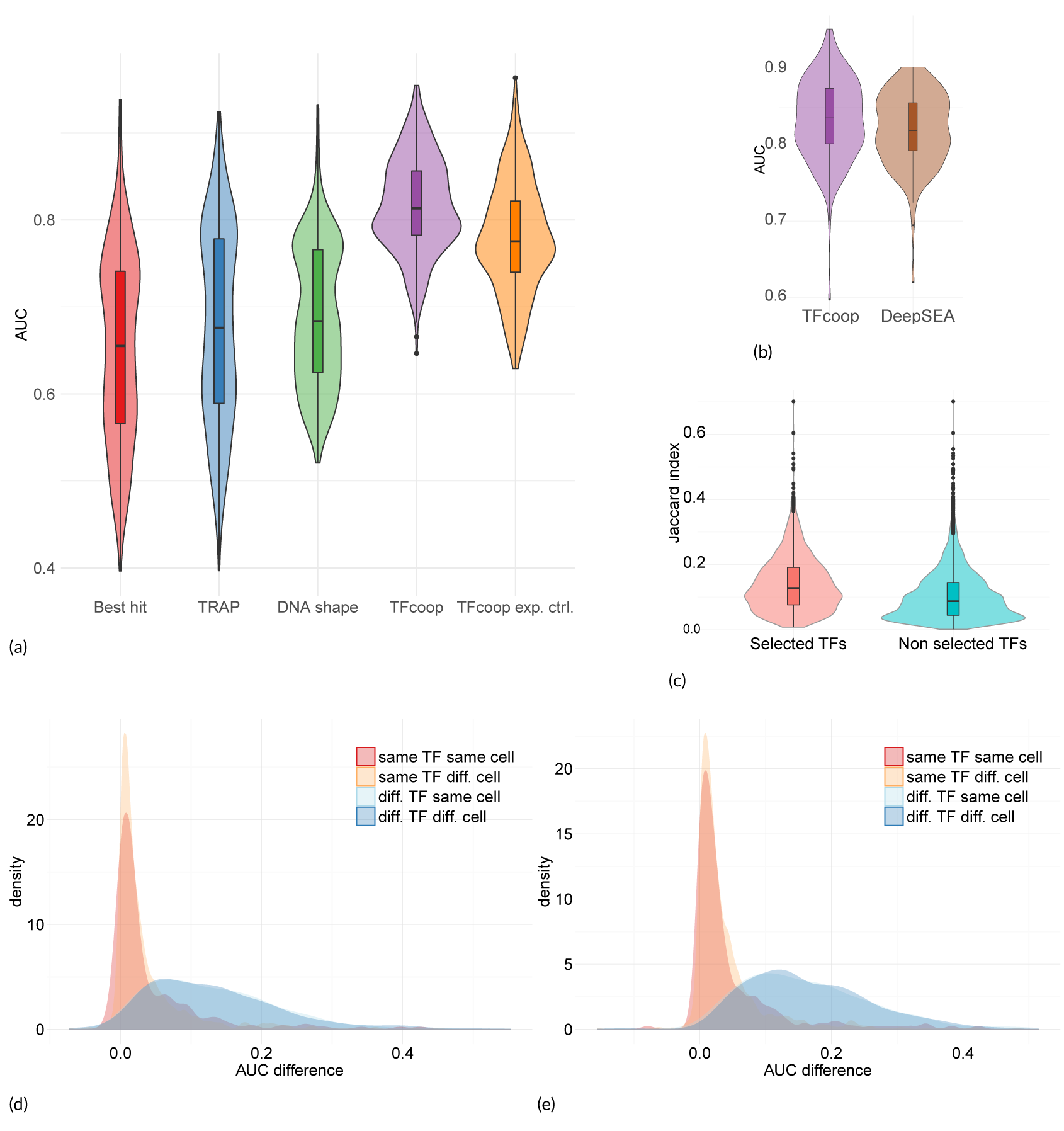
Accuracy and specificity on mRNA promoters. (a) Violin plots of the area under the ROC curves obtained in the 409 ChIP-seq. Best hit (red), TRAP (blue), DNAshape (green), TFcoop with no expression control (purple), and TFcoop with expression control (orange). ROC curves for Best hit, TRAP and DNAshape were computed in the non expression-controlled case. (b) Comparison of AUC achieved by TFcoop and DeepSea approach [44]. Comparison was done on 214 ChIP-seq experiments for which the DeepSea server provides predictions. (c) Intersection between pairs of ChIP-seq experiments associated with TFs identified as cooperating in promoters. These violin plots report the distribution of Jaccard indexes computed between different pairs of Chip-seq experiments. Left: for each TF A, we measured the Jaccard index between promoters bound by A and promoters bound by a TF B whose PWM has been selected in the TFcoop model learned for A (cases B = A were not considered). Right: for each TF A, we measured the Jaccard index between promoters bound by A and promoters bound by TFs whose PWMs have not been selected in the A model. (d-e) Distribution of AUC differences obtained when using a model learned on a first ChIP-seq experiment to predict the outcome of a second ChIP-seq experiment. Different pairs of ChIP-seq experiments were used: experiments on the same TF and same cell type (red), experiments on the same TF but different cell type (yellow), experiments on different TFs but same cell type (light blue), and experiments on different TFs and different cell types (blue). For each pair of ChIP-seq experiment A-B, we measured the difference between the AUC achieved on A using the model learned on A, and the AUC achieved on A using the model learned on B. AUC differences were measured on the non expression-controlled case (c) and on the expression-controlled case (d).

**FIGURE 2.**
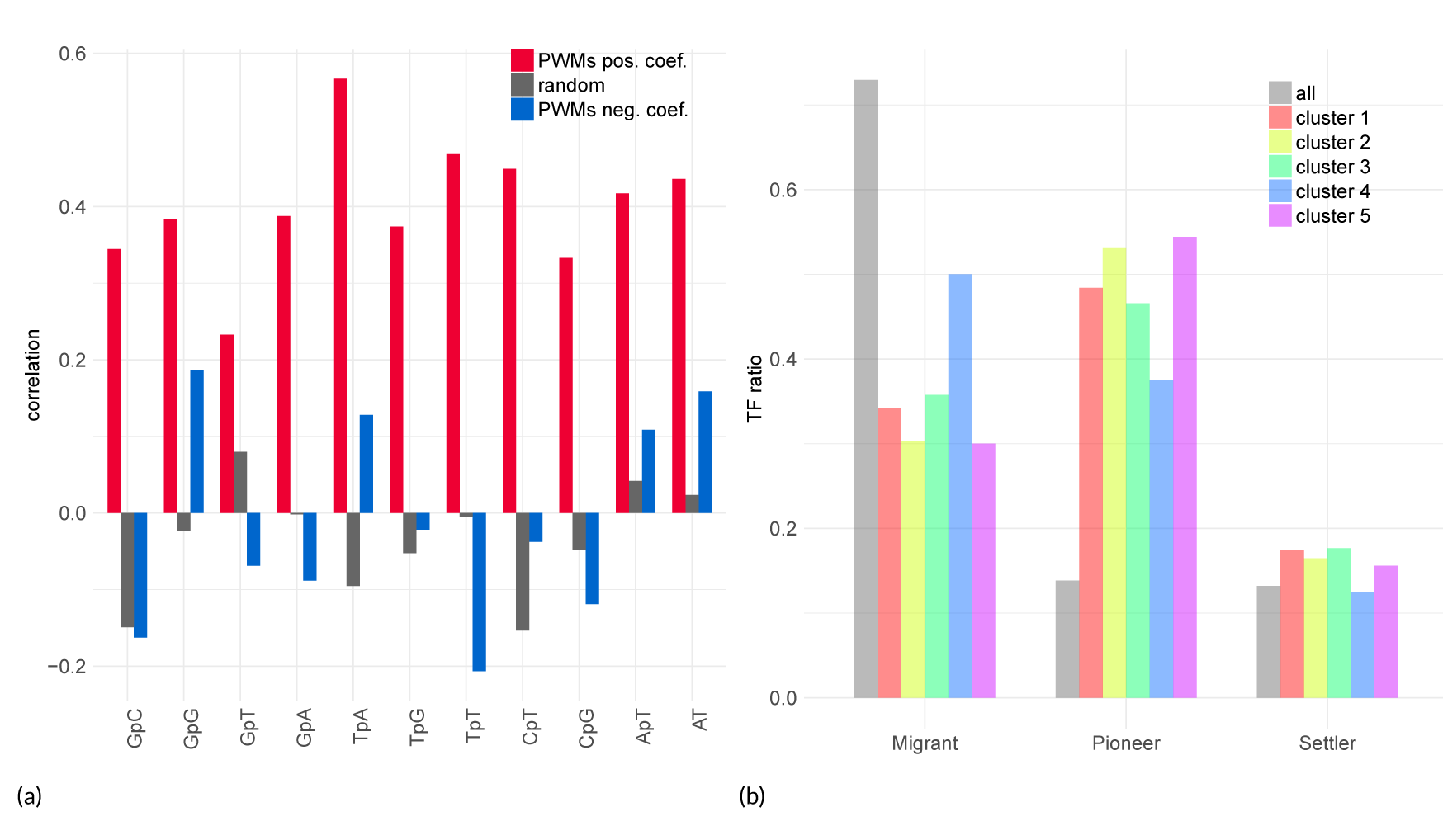
Selected PWMs in mRNA promoters. (a) Pearson correlation between nucleotide composition of the target PWM and the mean composition of selected PWMs (with positive and negative coefficients in red and blue, respectively) in 409 models. Grey: correlation achieved by randomly selecting the same number of PWMs for each model. (b) Pioneer TF distribution of selected PWMs in the different models. We kept one model for each target PWM to avoid bias due to over-representation of the same PWM in certain classes. Grey represents the distribution of all PWMs associated with a family in Sherwood et al. [9] (159 over 520 non-redundant PWMs)

**FIGURE 3.**
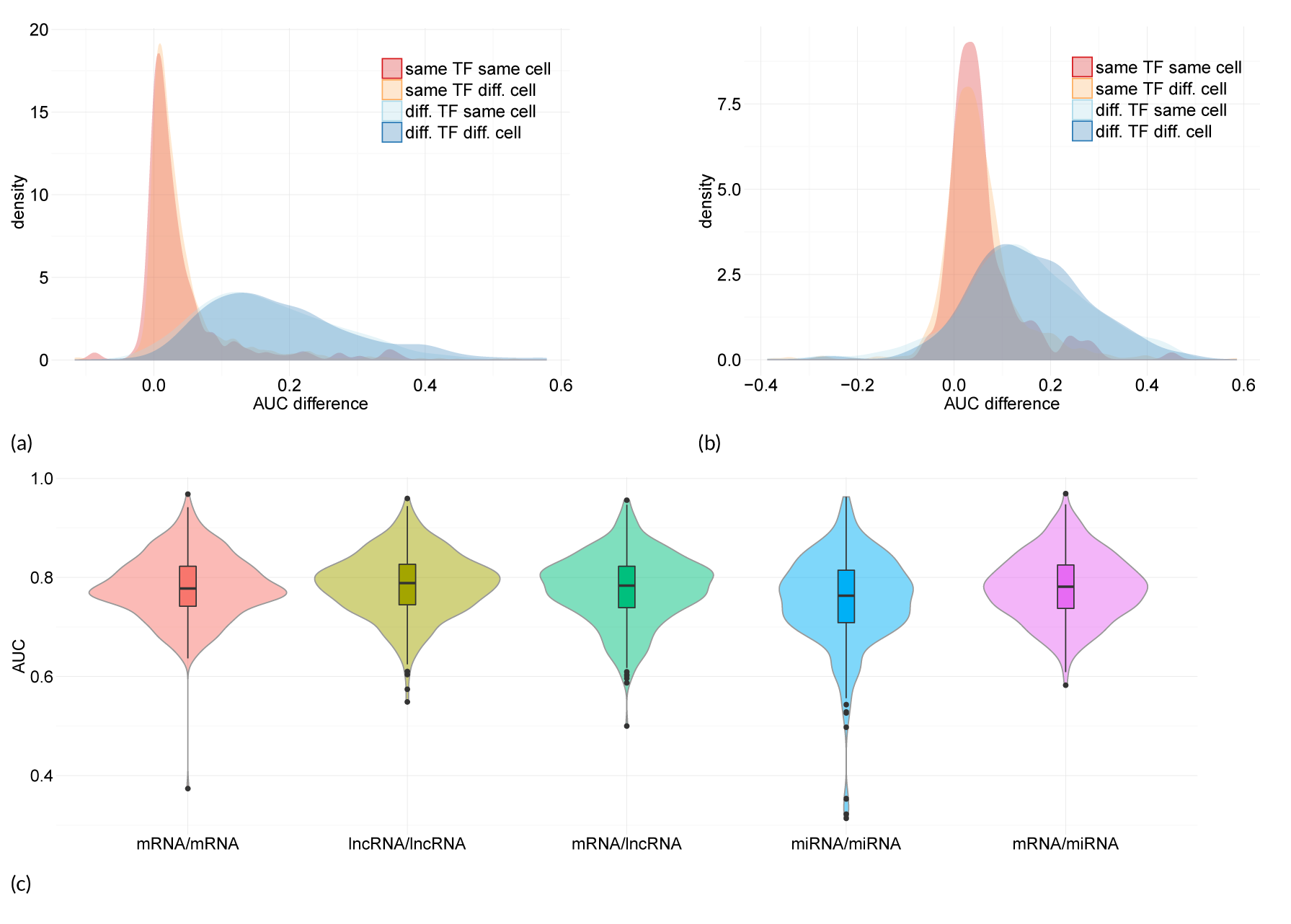
Accuracy and specificity on lncRNA and pri-miRNA promoters. Top: Model specificity on promoters of lncRNA (a) and pri-miRNAs (b). These figures represent the distribution of AUC differences obtained when using a model learned on a first ChIP-seq experiment to predict the outcome of a second ChIP-seq experiment. Different pairs of ChIP-seq experiments were used: experiments on the same TF and same cell type (red), experiments on the same TF but different cell type (yellow), experiments on different TFs but same cell type (light blue), and experiments on different TFs and different cell types (blue). For each pair of ChIP-seq experiment A-B, we measured the difference between the AUC achieved on A using the model learned on A, and the AUC achieved on A using the model learned on B. AUC differences were measured on the expression-controlled case. Bottom: Promoter models are interchangeable. For each ChIP-seq experiment, we computed the AUC of the model learned and applied on mRNAs (pink), learned and applied on lncRNAs (yellow-green), learned and applied on pri-miRNAs (blue), learned on mRNAs and applied to lncRNAs (green), learned on mRNAs and applied to pri-miRNAs (purple).

Next, we sought to assess the TF cooperations inferred by the models. If true, they should be apparent in the ChIP-seq experiments. Namely, if the PWM of TF B is among the selected variables for predicting the presence of TF A, then we should observe many common targets among the ChIP-seq experiments of TFs A and B. To test this, we first randomly selected one model for each different TF, and restricted our analyses to TFs associated with ENCODE ChIP-seq experiments. Then, for each model A, we measured the Jaccard index between promoters bound by TF A and promoters bound by a TF B whose PWM has been selected in model A (cases B = A were not considered), and we compared these scores to the same scores computed on TFs whose PWMs have not been selected in model A. A t-test score attests that Jaccard indexes are larger for selected TFs than for non-selected TFs (p-value < 1.*e*^-16^ and Figure 1(c)), and hence that the inferred TF cooperations are supported by experimental data.

Finally, we sought to take advantage of the relative redundancy of target TFs in the set of 409 ChIP-seq experiments to investigate the specificity of the learned models. Namely, we compared pairs of models learned from ChIP-seq experiments targeting (i) the same TF in the same cell-type, (ii) the same TF in different cell-types, (iii) different TFs in the same cell-type, and (iv) different TFs in different cell-types. In these analyses, we used the model learned on one ChIP-seq experiment A to predict the outcome of another ChIP-seq experiment B, and we compared the results to those obtained with the model directly learned on B. More precisely, we measured the difference of the Area under the ROC Curves (AUC) between the model learned on A and applied on B and the model learned and applied on B. To avoid any effect driven by the over-representation of CTCF in ChIP-seq data, we randomly selected only 10 ChIP-seq experiments targeting this TF in these analyses. As shown in Figure 1(d) and 1(e), models learned on the same TF (whether or not on the same cell-type) have overall smaller AUC differences than models learned on different TFs.

We then analyzed cell and TF specificity more precisely. Cell specificity refers to the ability of a model learned on one TF and in one cell type to predict the outcome of the same TF in another cell type. Oppositely, TF specificity refers to the ability of a model learned on one TF in one cell type to predict the outcome of another TF in the same cell type. Cell and TF specificities were evaluated by the shift between the associated distributions of AUC differences in Figure 1(d): cell specificity was assessed by the shift between red and yellow distributions, while TF specificity was assessed by the shift between red and light blue distributions. We used a standard t-test to measure that shift. Low p-values indicate high distribution shifts (hence high cell/TF specificity), while high p-values indicate low shifts (hence low specificity). Our results indicate very low cell specificity (p-values 0.91 and 0.95 in the non-controlled and expression-controlled cases, respectively) and high TF specificity (1 · 10^−61^ and 3 · 10^−83^). The fact that the TF specificity is slightly higher in the expression-controlled case suggests that part of the TF combinations that help discriminate between bound and unbound sequences is common to several TFs in the non-controlled case. It is indeed known that the majority of ChIP-seq peaks are found in open and active promoters [39]. Thus, most positive examples are associated with open chromatin marks. However, in the non-expression-controlled case, a large part of the negative examples are in closed chromatin and are therefore likely associated with other chromatin marks. As a result, in this case, TFcoop presumably also learns the TFBS signature that helps differentiate between these chromatin marks. Oppositely, in the expression-controlled case, the positive and negative examples have similar chromatin states, and TFcoop unveils the TFBS signature specific to the target TF. We can also observe that this renders the former problem slightly easier than the second one, as illustrated by the difference of TFcoop performances in Figure 1(a). Finally the low cell specificity means that the general rules governing TFBS combination in promoters do not dramatically change from one tissue to another. This is important in practice because it enables us to use a model learned on a specific ChIP-seq experiment to predict TBFSs of the same TF in another cell type.

### Analysis of TFBS combinations in promoters

We next analyzed the different variables (PWM scores and (di)nucleotide frequencies) that were selected in the 409 learned models. Overall, 95% of the variables correspond to PWM scores. Although only 5% of the selected variables are (di)nucleotide frequencies, almost all models include at least one of these features. As mentioned earlier, the learning algorithm does not use any prior knowledge and can select the variables that best help predict the ChIP-seq experiment without necessarily selecting the PWM of the target TF. Our analysis shows that, for 75% of the models, at least one version of the target PWM was selected. Moreover, it is important to note that similar PWMs tend to have correlated scores. Hence, another PWM may be selected instead of the target. To overcome this bias, we also considered all PWMs similar to the target PWM. We used Pearson correlation between PWM scores in all promoters to measure similarity and set a threshold value of 0.75 to define the list of similar PWMs. With this threshold, 90% models include the target or a similar PWM. Analysis of the remaining 10% models shows that they often correspond to ChIP-seq experiments with low number of positive sequences (median number 955 *vs.* 2477 for all ChIP-seq experiments). This may be due either to technical problems, to lowly expressed TFs, or to TFs that rarely bind promoters.

We further analyzed the most selected PWMs. To avoid any bias linked to the number of CTCF ChIP-seq experi-ments, we only considered 10 CTCF models that were randomly selected for the analyses. We ranked the PWMs by the number of models in which they appear, and look for enrichment of certain JASPAR structural families (bHLH, Zinc finger). A gene set enrichment analysis (GSEA) [52] shows that “tryptophan cluster factors” (FDR q-val< 10^−4^), “C2H2 zinc finger factors” (FDR q-val< 10^−4^) and “basic leucine zipper factors” (FDR q-val= 2 · 10^−3^) are the most represented classes of PWMs selected in the models (Supp. Figure 4). We then looked at the differences between models learned in the expression-controlled experiments and models learned in the non-controlled experiments. For each non-controlled model, we enumerated the variables that are selected in this model and not selected in the corresponding expression-controlled model. Several PWMs are over-represented in this list (see Supp. Table 1). Specifically, a GSEA analysis shows that the FOX family is particularly enriched (FDR q-val 7 · 10^−3^). Because FOXA is a well known pioneer factor [53], this enrichment is in agreement with our idea that, in the non-controlled case as opposed to the expression-controlled case, TFcoop also learns TFBS signatures linked to active/inactive chromatin marks.

**FIGURE 4.**
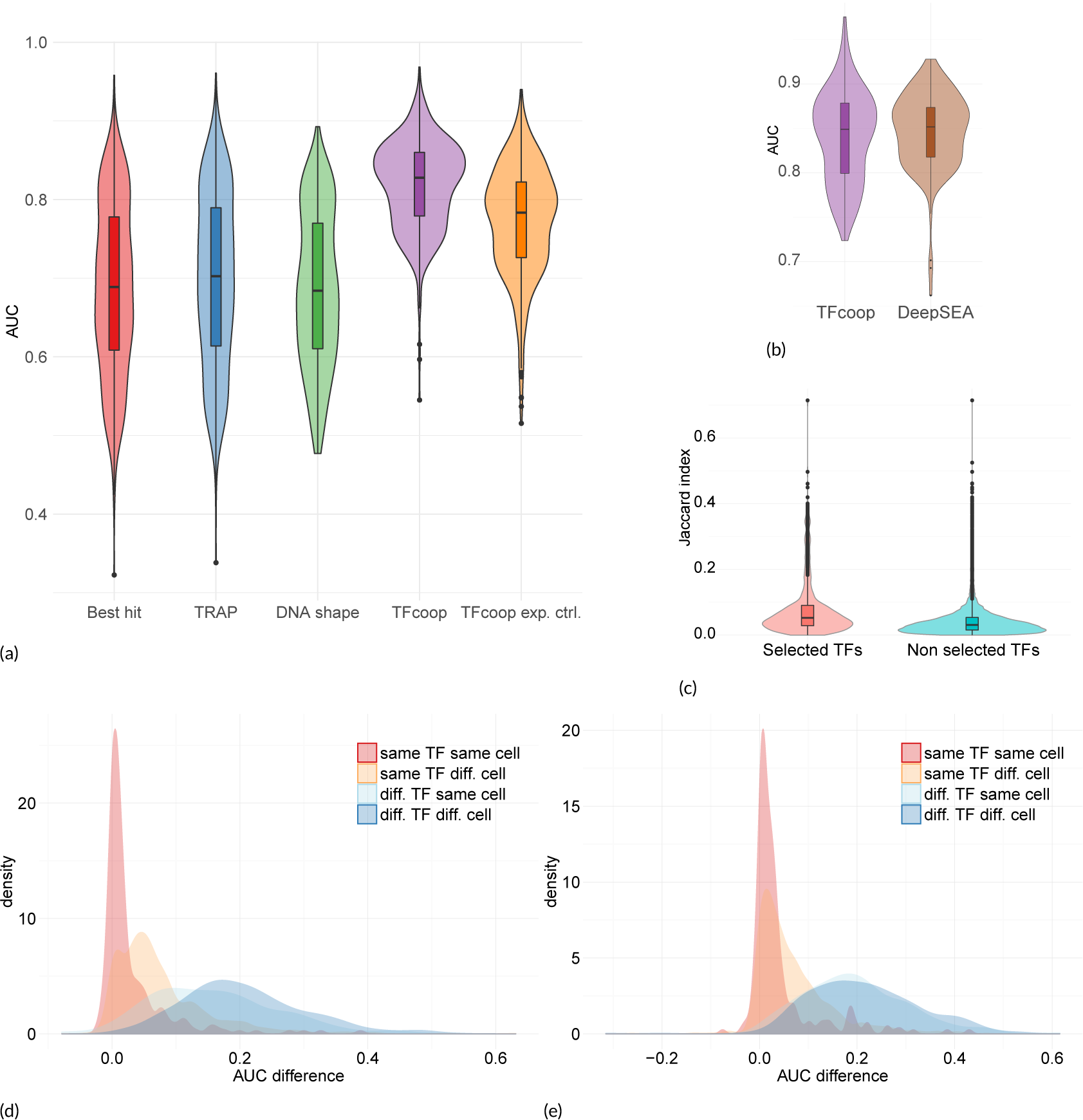
Accuracy and specificity on enhancers. (a) Violin plots of the area under the ROC curves obtained in the 409 ChIP-seq. Best hit (red), TRAP (blue), DNAshape (green), TFcoop with no expression control (purple), and TFcoop with expression control (orange). ROC curves for Best hit, TRAP and DNAshape were computed in the non expression-controlled case. (b) Comparison of AUC achieved by TFcoop and DeepSea approach [44]. Comparison was done on 214 ChIP-seq experiments for which the DeepSea server provides predictions. (c) Intersection between pairs of ChIP-seq experiments associated with TFs identified as cooperating in enhancers. These violin plots report the distribution of Jaccard indexes computed between different pairs of Chip-seq experiments. Left: for each TF A, we measured the Jaccard index between enhancers bound by A and enhancers bound by a TF B whose PWM has been selected in the TFcoop model learned for A (cases B = A were not considered). Right: for each TF A, we measured the Jaccard index between enhancers bound by A and enhancers bound by TFs whose PWMs have not been selected in the A model. (d-e) Distribution of AUC differences obtained when using a model learned on a first ChIP-seq experiment to predict the outcome of a second ChIP-seq experiment on enhancers. Different pairs of ChIP-seq experiments were used: experiments on the same TF and same cell type (red), experiments on the same TF but different cell type (yellow), experiments on different TFs but same cell type (light blue), and experiments on different TFs and different cell types (blue). For each pair of ChIP-seq experiment A-B, we measured the difference between the AUC achieved on A using the model learned on A, and the AUC achieved on A using the model learned on B. AUC differences were measured on the non expression-controlled case (b) and on the expression-controlled case (c).

**FIGURE 5.**
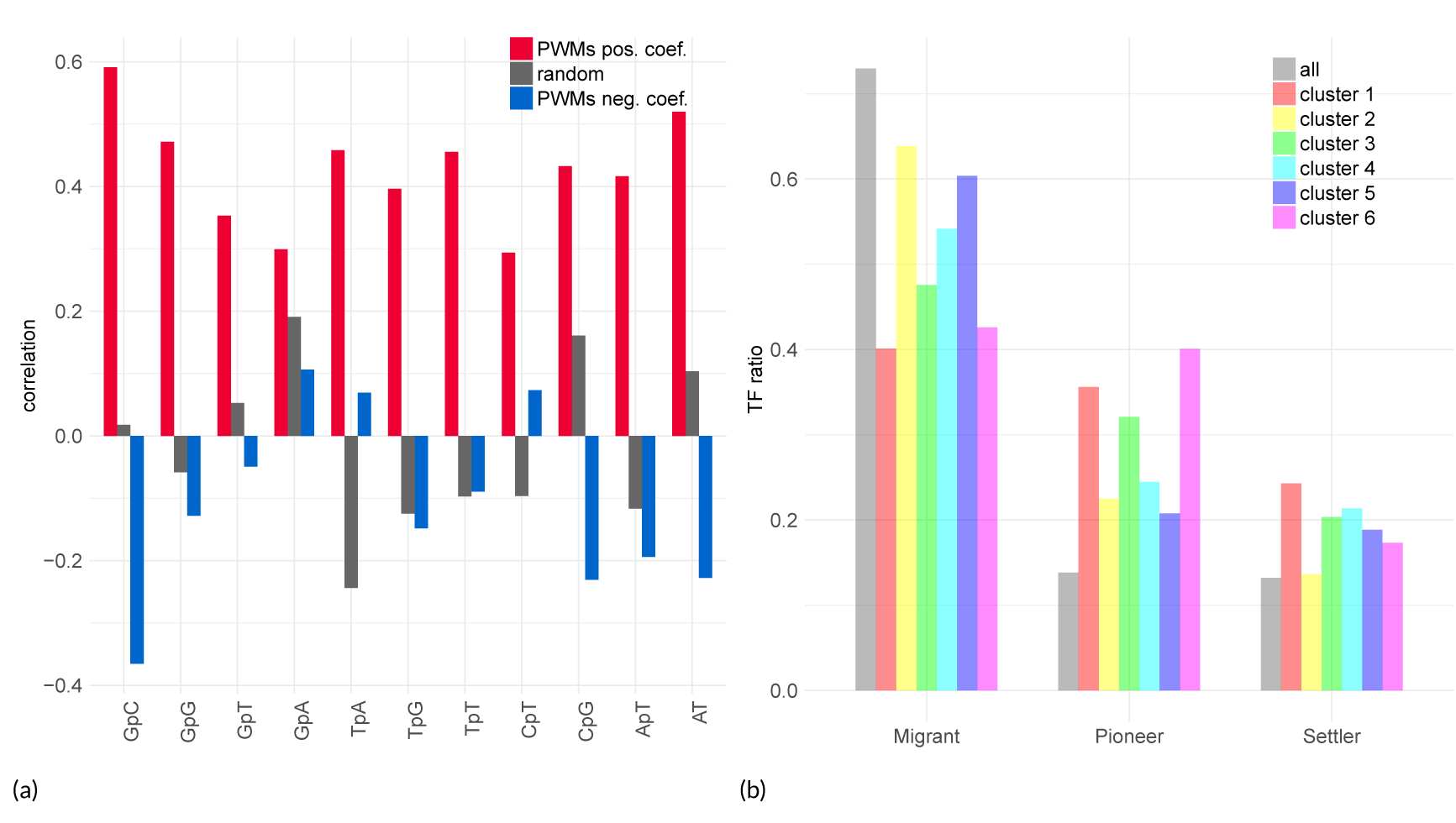
Selected PWMs in enhancers. (a) Pearson correlation between nucleotide composition of the target PWM and the mean composition of selected PWMs (see legend of Figure 2(a)) (b) Pioneer TF distribution in selected PWMs (same legend as Figure 2(b)).

Next, following the analyses of Levo et al. [32] and Dror et al. [33] we used our models to investigate the link between the nucleotide composition of the target PWM and that of the TFBS flanking region. First, we did not observe a significant link between target PWM composition and the (di)nucleotide variables that were selected in the models (Kolmogorov-Smirnov test p-val=0.448; see Supp. Figure 6). However, the (di)nucleotide composition of target PWM exhibited strong resemblance to that of the other selected PWMs (see Figure 2(a)). Specifically, the nucleotide and dinucleotide frequencies of the target PWM were strongly positively correlated with that of the PWMs selected with a positive coefficient. For PWMs selected with a negative coeficient the correlations are moderate or negative. This is in accordance with the findings of Dror et al. [33], who show that TFBS flanking regions often have similar nucleotide composition as the the TFBS.

**FIGURE 6.**
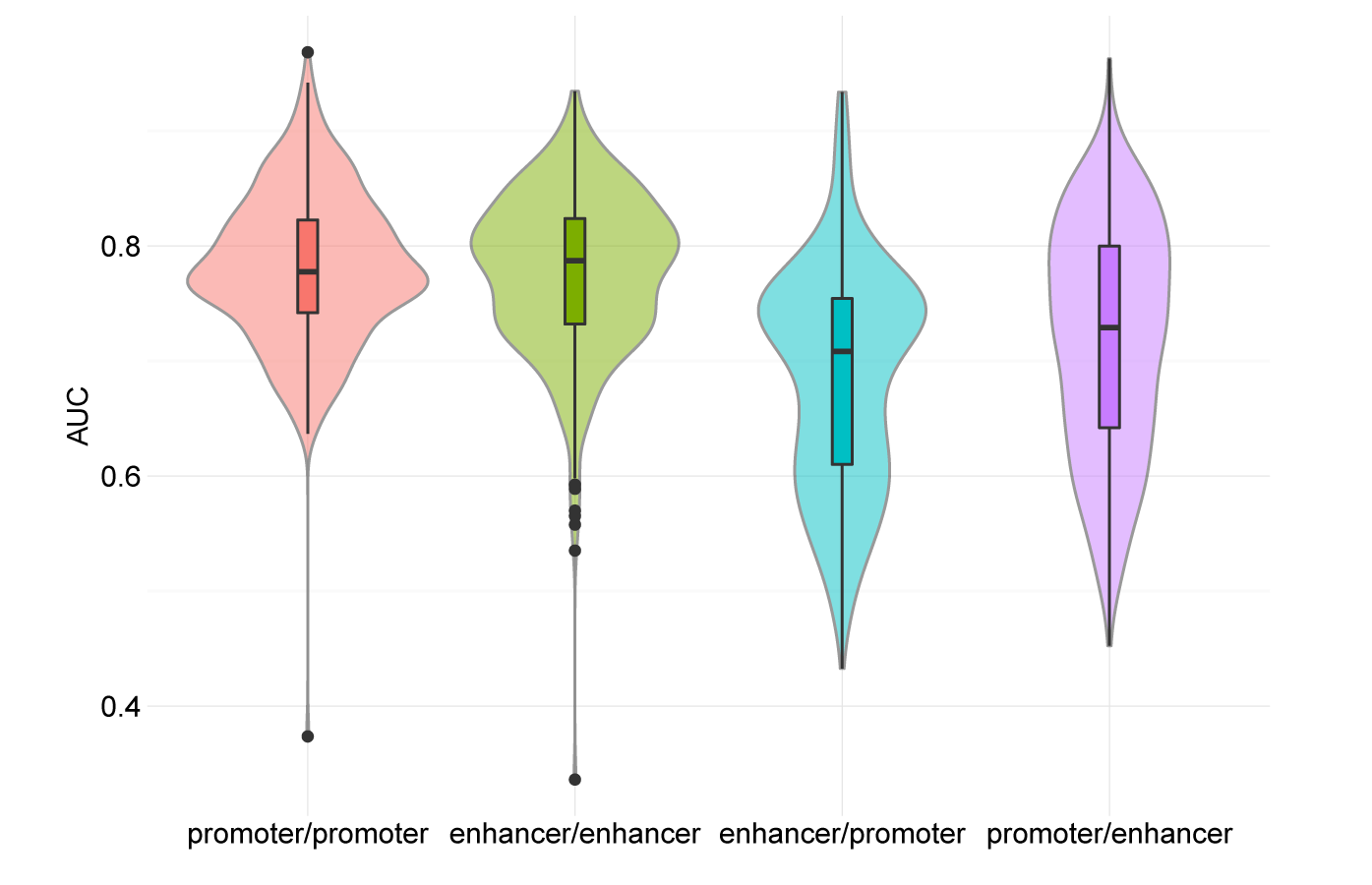
AUCs obtained in mRNA promoter and enhancer models. For each ChIP-seq experiment we computed the AUC of the model learned and applied on the promoters (red), learned and applied on the enhancers (green), learned on enhancers and applied to promoters (blue), and learned on promoters and applied to enhancers (purple).

We next evaluated the possibility of clustering the 409 learned models using the selected variables. As shown in Supp. Figure 7, the models can be partitioned in a few different classes with a k-means algorithm (5 classes were used in this figure). Supp. Figure 8 reports the most used variables in these different classes. We can first observe that, in agreement with our analysis of model specificity, the models associated with the same TF tend to cluster together. For example, the 4^*t*^ ^*h*^ class of our clustering is exclusively composed of CTCF models. Note that we did not observe any enrichment for the classical TF structural families (bHLH, Zinc finger,) in the different classes (data not shown). Actually, the clustering seems to be essentially driven by the nucleotide composition of the PWMs belonging to the models (see Supp. Figure 9).

**FIGURE 7.**
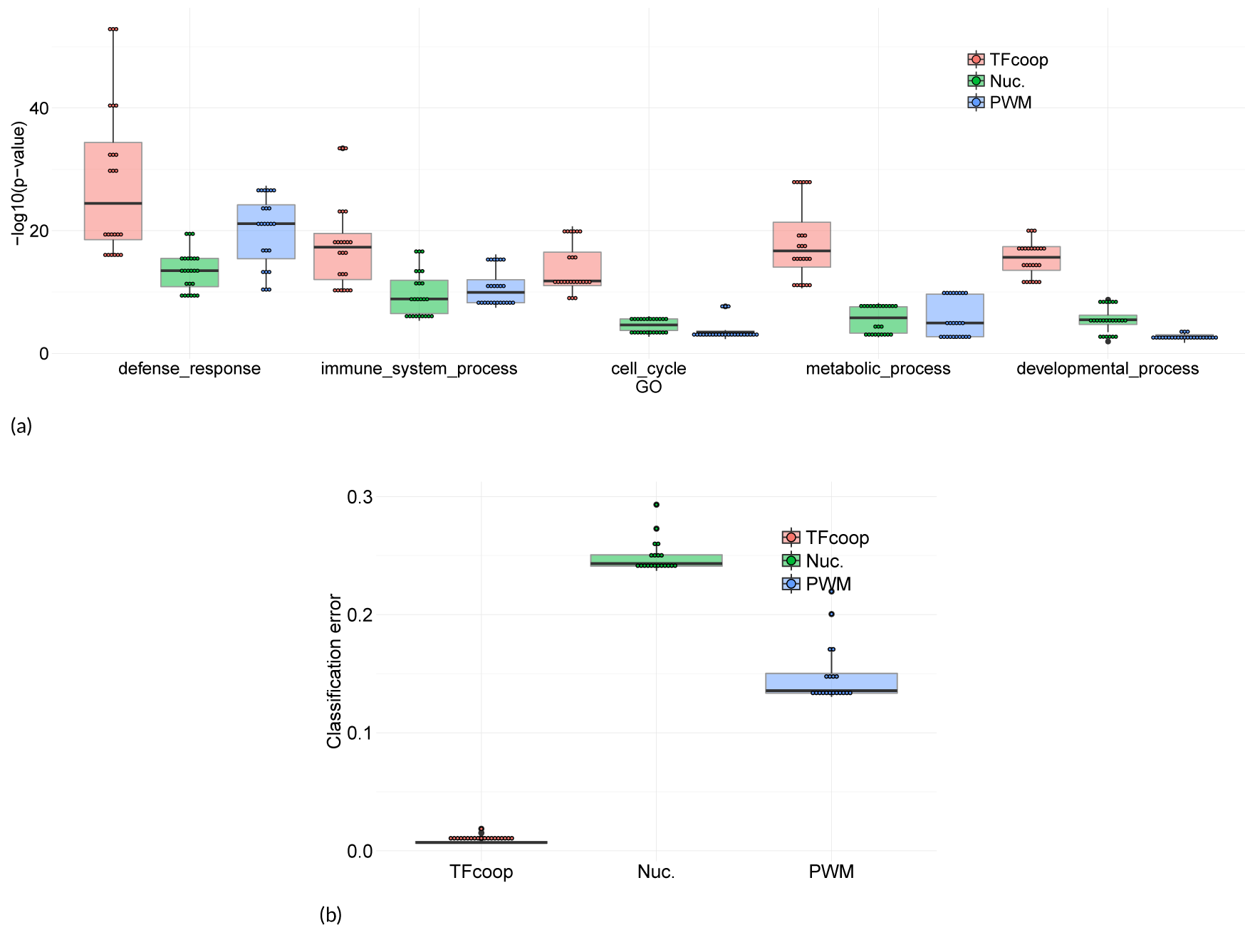
Using TFcoop scores for describing regulatory sequences. (a) GO term enrichment obtained with different promoter descriptions. Promoters were described using three different representations—TFcoop scores (red), (di)nucleotide frequencies (green), classical PWM scores (blue)— and then partitioned several times with different eans and different class numbers (see main text). For each clustering we identified the best p-value (Fisher exact test) associated with 5 GO terms (“defense response”, “immune system process”, “cell cycle”, “metabolic process”, “developmental process”) in any cluster. (b) Classification errors achieved with KNN classifiers discriminating between promoter and enhancer sequences. Boxplots describe the errors obtained using TFcoop scores (red), (di)nucleotide frequencies (green), and the classical PWM scores (blue), using different number of neighbors (K).

Pioneer TFs are thought to play an important role in transcription by binding to condensed chromatin and enhancing the recruitment of other TFs [9]. As shown in Figure 2(b) and by a GSEA analysis (Supp. Figure 5), pioneer factors clearly are over-represented in the selected variables of the models, whereas they represent less than 14% of all TFs. These findings are in agreement with their activity: pioneer TFs occupy previously closed chromatin and, once bound, allow other TFs to bind nearby [9]. Hence the binding of a given TF requires the prior binding of at least one pioneer TF. We also observed that TFs whose binding is weakened by methylation [50] are enriched in all models (Supp. Figure 10). This result may explain how CpG methylation can negatively regulate the binding of a given TF *in vivo* while methylation of its specific binding site has a neutral or positive effect *in vitro* [50]: regardless of the methylation status on its binding site, the binding of a TF can also be influenced *in vivo* by the sensitivity of its partners to CpG methylation.

### TFBS combinations in lncRNA and pri-miRNA promoters

We then ran the same analyses on the promoters of lncRNAs and pri-miRNAs using the same set of ChIP-seq experi-ments. Results are globally consistent with what we observed on mRNA promoters (see Figure 3 for the expression-controlled case). Overall, models show good accuracy and specificity on lncRNAs. Models are less accurate and have lower specificity for pri-miRNAs but this likely results from the very low number of positive examples available for these genes in each ChIP-seq experiment (Supp. Figure 11), which impedes both the learning of the models and estimation of their accuracy.

Next we sought to compare the models learned on mRNA promoters to the models learned on lncRNA and pri-miRNA promoters. For this, we interchanged the models learned on the same ChIP-seq experiment, *i.e.* we used the model learned on mRNA promoters to predict the outcome on lncRNA and pri-miRNA promoters. One striking fact illustrated by Figure 3(c) is that models learned on mRNA promoters and those learned on lncRNA promoters are almost perfectly interchangeable. This means that the TFBS rules governing the binding of a specific TF in a promoter are similar for both types of genes. We obtained consistent results when we used the models learned on mRNAs to predict the ChIP-seq outcomes on pri-miRNA promoters (Figure 3(c)). Accuracy is even better than that obtained by models directly learned on pri-miRNA promoters, illustrating the fact that the poor performance achieved on pri-miRNA promoters likely results from the small number of learning examples available for these genes.

### TFBS combinations in enhancers

We next applied the same approach on 38,554 enhancers defined by the FANTOM consortium [49]. We used the same ChIP-seq experiments as for the promoters. All enhancer sequences overlapping a ChIP-seq peak in the considered ChIP-seq experiment were considered as positive examples. As for promoters, we used two strategies to select positives and negative examples: in a first case we did not apply any control on their expression, while in a second case, we used CAGE expression data in the different tissues to only select expressed enhancers.

As observed for promoters, TFcoop outperforms classical PWM-based approaches on many TFs (see Figure 4(a) and Supp. Figure 12) and achieves results close to that of DeepSea [44] (Figure 4(b)). Here again, the non expression-controlled problem seems slightly easier than the controlled one. Using the same “Jaccard index test” used for promoters,we also assessed that the TF cooperations inferred by the models can be observed in ChIP-seq data and hence are likely to be biologically valid (p-value < 1.*e*^-16^ and Figure 4(c)).

However, analysis of model specificity reveals somewhat different results from that observed for promoters. Globally, models have good TF specificity: models learned on the same TF have more similar prediction accuracy than models learned on different TFs. However, in contrast to promoters, cell specificity is high in the non-controlled case (p-value 2·10^−45^; see peak shift in Figure 4(d)), although much lower in the expression-controlled case (p-value 1.·10^12^). Additionally, TF specificity seems slightly higher in the expression-controlled case than in the non-controlled case (p-values 1.7·10^−102^ *vs.* 1.·10^−114^). This is in accordance with our hypothesis formulated for promoters, that part of the TF combinations learned by TFcoop in the non-controlled case actually differentiates between active and inactive chromatin marks. This also seems to indicate that these TF combinations are cell-type specific, while the remaining combinations are more general (as illustrated by the 1.6 ·10^−12^ p-value measured on the expression-controlled case). Moreover, analysis of selected variables reveals that models learned without expression control involve much more variables than models learned with expression control (median numbers 18 *vs.* 11; t-test p-value ∼ 10^−9^). As a consequence, several variables are statistically more abundant in non-controlled models than in the cognate expression-controlled models (see Supp. Table 1). Interestingly, among the four variables with the most important differences, three are dinucleotides CpG, TpC and ApT. This may indicate that part of the active/inactive chromatin marks is linked to the dinucleotide composition of the underlying sequence. This proposal is in line with findings revealing the existence of sequence-level instructions for chromatin modifications[44, 45, 54]. Moreover, a GSEA analysis shows that the PWMs with the strongest differential enrichments belong to the “three-zinc finger kruppel-related factors” ((FDR q-val 1 · 10^−2^)). As some of these factors, in particular KLF1 [55], are linked to chromatin remodeling, this enrichment supports the idea that TFcoop also identifies TF combinations linked to epigenetics. The fact that cell-type specificity is more apparent for enhancers than for promoters in the non expression-controlled case (2 · 10^−45^ for enhancers *vs.* 0.91 for promoters) is in accordance with the fact that, contrary to promoters, most enhancers are expressed in a cell-specific manner (Supp. Figure 13 and ref. [49]).

As for promoters, we observed that the selected PWMs tends to have similar (di)nucleotide composition as the target PWM (Figure. 5(a)). Moreover, models can also be partitioned in a few different classes according to the selected variables (Supp. Figures 14 and 15). These classes mostly correspond to the nucleotide composition of the target and selected PWMs (Supp. Figure 16). Pioneer TFs are also over-represented in the selected PWMs (Figure 5(b) and Supp. Figure 5).

Next we sought to compare the models learned on enhancers to the models learned on promoters. First, we observed that enhancer models involve PWMs that are different from that used in promoter models (Supp. Table 2). Note for instance that several AP-1 TFs (FOS/JUN) are enriched in enhancers, in accordance with their prominent role in enhancer selection [56]. The same three structural classes are found enriched, but in different proportions, with more “C2H2 zinc finger factors” in promoters and more “basic leucine zipper factors” in enhancers (Supp. Figure 4). In term of prediction, promoter and enhancer models have globally similar accuracy (see Figure 6 on the expression-controled cases). However, a pairwise comparison of the models learned on each ChIP-seq experiment shows that the prediction accuracy is only moderately correlated (Pearson correlation 0.33; see Supp. Figure 17). Moreover, if we interchange the two models learned on the same ChIP-seq experiment, we observe that the model learned on promoters is generally not as good on enhancers as it is on promoters and *vice-versa* (Figure 6). Hence, while the rules learned on enhancers (promoters) in a given cell type are globally valid for enhancers (promoters) of other cell types, they do not apply to promoters (enhancers) of the same cell type. Note that AUCs of models learned on promoters and applied to enhancers are greater than that of models learned on enhancers and applied to promoters (Figure 6). This result might be explained by the existence of promoters able to exert enhancer functions [57, 58]. Conversely, the FANTOM definition of enhancers precludes potential promoter functions [49].

### Using TFcoop scores to describe regulatory sequences

We next explored whether TFcoop scores could be used to provide meaningful descriptions of regulatory sequences. This was assessed in two ways. First, we used the TFcoop models to cluster mRNA promoters and searched for over-represented gene ontology (GO) terms in the inferred clusters. We randomly selected one model for each TF, and used the 106 selected models to score the 20,846 mRNA promoter sequences. Each promoter sequence was then described by a vector of length 106. We next ran a k-means algorithm to partition the promoters into 5 different clusters, and we searched for over-represented GO terms in each cluster. For comparison, we ran the same procedure using two other ways to describe the promoter sequences: the classical PWM scores of the same 106 selected TFs (so promoters are also described by vectors of length 106), and the (di)nucleotide frequencies of the promoters (vector of length 12). Globally, the same GO terms appear to be over-represented in the different gene clusters and the three different clusterings: defense response, immune system process, cell cycle, metabolic process, and developmental process. We noticed that the p-values obtained with the TFcoop scores were invariably better than the two others. To avoid any clustering bias, we repeated the k-means clusterings several times, with various numbers of clusters. Namely, for each approach we ran 3 clusterings for each number of clusters ranging between 3 and 10 (resulting in 24 different clusterings for each approach) and computed over-representation p-values for the 5 GO terms in each cluster. As shown in Figure 7(a), the TFcoop scores substantially and systematically outperform the other scoring functions, indicating that the classification obtained with this score is more accurate to functionally annotate promoters than the others. Implicitly, these results are also consistent with a model in which most biological processes are controlled by specific combinations of TFs.

Next, we used the TFcoop models to discriminate between mRNA promoters and enhancers. We randomly split the sets of promoters and enhancers in training and test sets, and learned a K-nearest neighbor (KNN) classifier for discriminating between promoter and enhancer sequences on the basis of scores of the TFcoop models learned on promoters. As above, we also used the classical PWM scores of the same 106 selected TFs and (di)nucleotide frequencies of the sequences. We resumed the procedure with a number of neighbors (K) varying between 1 and 20, and computed the number of errors obtained by each approach on the test set (Figure 7(b)). Here again, TFcoop description outperforms other description methods, with an error rate around 2% for TFcoop *vs.* 15% and 25% for the other approaches. This result confirms the existence of DNA features distinguishing enhancers from mRNA promoters [49, 21] and identifies TF combinations as potent classifiers.

### Identifying TFs responsible for gene expression change

As a final test, we sought to use TFcoop to identify the TFs responsible for gene expression change in various gene expression experiments. For this, we used the compendium collected by Meng et al. [59]. The interest of this compendium is that each data corresponds to a particular TF for which the activity has been modified (repressed or enhanced), hence the TF responsible for deregulation (hereafter called as the “responsible TF”) is known. In each experiment, we selected the top 500 genes with the highest positive log fold change, and computed the difference of score distribution of the responsible TF in the top 500 promoters and in all other promoters with a Kolmogorov-Smirnov test. This was done using the classical PWM scoring function and with the TFcoop scores. Of the 21 experiments, 5 responsible TFs achieved enrichment p-values below 1% with the classical PWM scoring function, while this number rises to 13 with the TFcoop score (see Supp. Table 3).

One striking fact however is that numerous TFs (not solely the responsible TF) appear to be also enriched in the top 500 promoters (Supp. Table 3). Note that this effect is not restricted to the TFcoop scoring. The classical PWM scoring method also has numerous enriched TFs on the experiments for which it yields good p-values on the responsible TF. There can be different explanations for this effect. First, modifying the activity of the responsible TF may induce a cascade of activations/repressions of other TFs. Second, if two TFs A and B often bind together in promoters, they may share a high number of target genes. In this case, TF B may appear as over-represented in the promoters of genes deregulated by TF A, even if TF B is not itself deregulated. This provides us with an interesting way to assess our models. Namely, when this appends (and if our models are meaningful) then TF A should be present among the selected variables of the TF B model. For each experiment, we therefore enumerated all TFs enriched in the top 500 promoters and checked whether the responsible TF was present in their models. We used a Fisher exact test to assess whether this appends more often than expected in the different experiments (Supp. Table 3). Of the 18 testable experiments, 13 yield a p-value below 5%, indicating that the responsible TF is often involved in the TF combinations associated with the TFs enriched in the top 500 promoters.

## DISCUSSION

In this paper we proposed a method to identify TF combinations that can be predictive of the binding of a target TF. Our approach is based on a logistic model learned from ChIP-seq experiments on the target TF. Cross-validation study showed that the approach is effective and outperforms classical PWM based approaches on many TFs. It is important to note that TFcoop combinations do not necessarily reflect just cooperation, but also competition. For instance, a TF A competing with a TF B may be useful to predict the binding of B and would thus appear in the TF B model while A and B do not cooperate.

We distinguished two prediction problems associated with two situations, depending whether the aim is to predict binding in any promoter/enhancer or solely in expressed promoters/enhancers. For expressed promoters/enhancers, our experiments showed that the learned models have high TF specificity and quite low cell-type specificity. On the other hand, for the problem of expressed and not expressed promoters/enhancers binding, the learned models are less TF specific and more cell-type specific (especially for enhancers). These results are in accordance with a two-level model of gene regulation: (i) cell-type specific level that deposits specific chromatin marks on the genome, and (ii) non, or poorly, cell-type specific level that regulates TF binding in all DNA regions associated with appropriate marks.

An important property highlighted by our models is that rules governing TF combinations are very similar in the promoters of the three gene types analyzed (mRNA, pri-miRNA and lncRNA), but different between promoters and enhancers. This is further confirmed by our experiments for discriminating between promoter and enhancer sequences showing that scores produced by TFcoop models allow accurate classification between the two types of sequences. Our results thus argue for a prominent role of transcription factor binding as the fundamental determinant of regulatory activity able to distinguish enhancers and promoters [21]. Furthermore, as promoters and enhancers produce different RNA molecules [49, 21], our results also suggest that the production of enhancer RNAs (eRNAs) on one hand, and that of mRNAs, lncRNAs, and miRNAs on the other hand, requires a specific and distinct subset of TFs.

Our approach could be improved in several ways. A quite straightforward improvement would be to use the DNAshape score developed by Mathelier et al. [36] instead of the classical PWM score. This could improve TFcoop accuracy for several TFs, especially for TFs such as CTCF for which TFcoop does not outperform classical PWM scoring. More profoundly, one drawback of TFcoop is that the logistic model enables us to learn only a single TF combination for each target TF. However, we can imagine that certain TFs may be associated with two or more different TF combinations depending on the promoter/enhancer they bind. A solution for this would be to learn a discrimination function based on several logistic models instead of a single one.

## MATERIAL AND METHODS

### Promoter, enhancer, long non-coding RNA and microRNA sequences

We predicted TF binding in both human promoters and enhancers. For promoters, sequences spanning ± 500bp around starts (i.e. most upstream TSS) of protein-coding genes, long non-coding RNAs and microRNAs were considered. Starts of coding and lncRNA genes were obtained from the hg19 FANTOM CAGE Associated Transcriptome (CAT) annotation [48, 19]. Starts of microRNA genes (primary microRNAs, pri-miRNAs) were from [20]. For enhancers, sequences spanning ± 500bp around the mid-positions of FANTOM-defined enhancers [49] were used. Lastly, our sequence datasets are composed of 20,845 protein coding genes, 1,250 pri-microRNAs, 23,887 lncRNAs, and 38,553 enhancer sequences.

### Nucleotide and dinucleotide features

For each of these sequences, we computed nucleotide and dinucleotide relative frequencies as the occurrence number in the sequence divided by sequence length. Frequencies were computed in accordance with the rule of DNA reverse complement. For nucleotides, we computed the frequency of A/T and G/C. Similarly, frequencies of reverse complement dinucleotides (*e.g.* ApG and CpT) were computed together. This results in a total of 12 features (2 nucleotides and 10 dinucleotides).

### PWM

We used vertebrate TF PFMs from JASPAR [23], including all existing versions of each PFM, resulting in a set of 638 PFMs with 118 alternative versions. PFMs were transformed into PWMs as described in Wasserman and Sandelin [22]. PWM scores used by TFcoop for a given sequence were computed as described in [22], keeping the maximal score obtained in any position of the sequence. Namely, each PWM was used to scan the entire sequence and score each position, and the maximal score was used as potential predictive feature by TFcoop.

### ChIP-seq data

We collected ChIP-seq data from the ENCODE project [60] for human immortalized cell lines, tissues, and primary cells. Experiments were selected when the targeted TF were identified by a PWM in JASPAR. Thus we studied 409 ChIP-seq experiments for 106 distinct TFs and 41 different cell types. The most represented TF is CTCF with 69 experiments, while 88% of the experiments are designed from immortalized cell lines (mainly GM12878, HepG2 and K562). The detailed list of all used experiments is given in Supplementary materials. For each ChIP-seq experiment, regulatory sequences were classified as positive or negative for the corresponding ChIP targeted TF. We used Bedtools v2.25.0 [61] to detect intersection between ChIP-seq binding sites and regulatory sequences (both mapped to the hg19 genome). Each sequence that intersects at least one ChIP-seq binding region was classified as a positive sequence. The remaining sequences formed a negative set. The number of positive sequences varies greatly between experiments and sequence types. Mean and standard deviation numbers of positive sequences are respectively 2661(±1997) for mRNAs, 1699(±1151) for lncRNAs, 216(±176) for microRNAs, and 1516(±1214) for enhancers.

### Expression data

To control the effect of expression in our analyses, we used ENCODE CAGE data restricted to 41 cell lines. The expression per cell line was calculated as the mean of the expression observed in all corresponding replicates. For microRNAs, we used the small RNA-seq ENCODE expression data collected for 3,043 mature microRNAs in 37 cell lines (corresponding to 403 ChIP-seq experiments). The expression of microRNA genes (i.e. pri-microRNAs) was calculated as the sum of the expression of the corresponding mature microRNAs.

### Logistic model

We propose a logistic model to predict the regulatory sequences bound by a specific TF. Contrary to classical approaches, we not only consider the score of the PWM associated with the target TF, but also the scores of all other available PWMs. The main idea behind this is to unveil the TF interactions required for effective binding of the target TF. We also integrate in our model the nucleotide and dinucleotide compositions of the sequences, as the environment of TFBSs are thought to play major role in binding affinity [32, 33].

For each ChIP-Seq experiment, we learn different models to predict sequences bound by the target TF in four regulatory regions (promoters of mRNA, lncRNA and pri-miRNA, and enhancers). For a given experiment and regulatory region, our model aims to predict response variable *y*_*s*_ by the linear expression

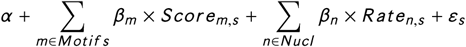

where *y*_*s*_ is the Boolean response variable representing the TF binding on the given sequence *s* (*y*_*s*_ = 1 for TF binding, 0 otherwise); *Scor e*_*m,s*_ is the score of motif *m* on sequence *s*; *Rate*_*n,s*_ is the frequency of (di)nucleotide *n* in sequence *s*; *α* is a constant; *β*_*m*_ and *β*_*n*_ are the regression coefficient associated with motif *m* and (di)nucleotide *n*, respectively;and *ε*_*s*_ is the error associated with sequence *s*. *M ot i f s* and *Nucl* sets respectively contain 638 JASPAR PWMs and 12 (di)nucleotide frequencies.

To perform variable selection (*i.e.* identifying cooperating TFs), we used the LASSO regression minimising the prediction error within a constraint over *l* 1-norm of *β* [47]. The weight of the LASSO penalty is chosen by cross-validation by minimising the prediction error with the R package *g l mnet* [62]. As the response variable is Boolean, we used a logistic regression giving an estimation of the probability to be bound for each sequence. We evaluate the performance of the model using 10-fold cross validation. In each validation loop, 90% of sequences (training data) are used to learn the *β* parameters and the remaining 10% (test data) are used to evaluate the predictive performance of the model.

## Alternative approaches

We compared the predictive accuracy of our model to three other approaches.

### Best hit approach

The traditional way to identify TF binding sites consists in scanning a sequence and scoring the corresponding PWM at each position. Positions with a score above a predefined threshold are considered as potential TFBS. A sequence is then considered as bound if it contains at least one potential TFBS.

### TRAP score

An alternative approach proposed by Roider et al. [51] is based on a biophysically inspired model that estimates the number of bound TF molecules for a given sequence. In this model, the whole sequence is considered to define a global affinity measure, which enables us to detect low affinity bindings. We use the R package *tRap* [62] to compute the affinity score of the 638 PWMs for all sequences. As proposed in [51], we use default values for the two parameters (*R*_0_(*width*), *λ* = 0.7).

### DNA shape

In addition to PWMs, Mathelier et al. [36] considered 4 DNA shapes to increase binding site identification: helix twist, minor groove width, propeller twist, and DNA roll. The 2^*nd*^ order values of these DNA shapes are also used to capture dependencies between adjacent positions. Thus, each sequence is characterized by the best hit score for a given PWM plus the 1^*st*^ and 2^*nd*^ DNA shape order values at the best hit position. The approach based on gradient boosting classifier requires a first training step with foreground (bound) and background (unbound) sequences to learn classification rules. Then the classifier is applied to the set of test sequences. We used the same 10-fold cross-validation scheme that we used in our approach. We applied two modifications to speed-up the method, which was designed for smaller sequences. First, in the PWM optimization step of the training phase, we reduced the sequences to ± 50bp around the position with highest ChIP-Seq peak for positive sequences and to ± 50bp around a random position for negative sequences. Second, after this first step we also reduced sequences used to train and test the classifiers to ± 50bp around the position for which the (optimized) PWM gets the best score.

### DeepSEA

Zhou and Troyanskaya [44] proposed a deep learning approach for predicting the binding of chromatin proteins and histone marks from DNA sequences with single-nucleotide sensitivity. Their deep convolutional network takes 1000bp genomic sequences as input and predicts the states associated with several chromatin marks in different tissues. We used the predictions provided by DeepSEA server (http://deepsea.princeton.edu/). Namely, coordinates of the analyzed promoter and enhancer sequences were provided to the server, and the predictions associated with each sequence were retrieved. Only the predictions related to the ChIP-seq data we used in our analyses were considered (*i.e.* 214 ChIP-seq data in total).

## ACKNOLEDGEMENTS

We thank Anthony Mathelier and Wyeth Wasserman for insightful discussions and suggestions. We are indebted to researchers around the globe who generated experimental data and made them freely available. This work was supported by funding from CNRS, *Plan d’Investissement d’Avenir* #ANR-11-BINF-0002 *Institut de Biologie Computationnelle* (young investigator grant to C-H.L. and post-doctoral fellowship to J.V.), Labex NUMEV (post-doctoral fellowship to J.V.), INSERM-ITMO Cancer project “LIONS” BIO2015-04, and CNRS International Associated Laboratory “miREGEN”.

## SUPPORTING INFORMATION

The R code for learning a TFcoop model from a ChIP-seq experiment is available in an R Markdown file at address https://gite.lirmm.fr/brehelin/TFcoop This file also provides the R scripts for reproducing the main experiments described in the paper. The different models learned on mRNA, lncRNA, miRNA promoters and enhancers are available as R object (.RData) at the same address.

